# A genetic probe for visualizing glutamatergic synapses and vesicles by 3D electron microscopy

**DOI:** 10.1101/2020.07.31.230995

**Authors:** Thomas Steinkellner, Matthew Madany, Matthias G. Haberl, Vivien Zell, Carolina Li, Junru Hu, Mason Mackey, Ranjan Ramachandra, Stephen Adams, Mark H. Ellisman, Thomas Hnasko, Daniela Boassa

## Abstract

Communication between neurons relies on the release of diverse neurotransmitters, which represent a key-defining feature of a neuron’s chemical and functional identity. Neurotransmitters are packaged into vesicles by specific vesicular transporters. However, tools for labeling and imaging synapses and synaptic vesicles based on their neurochemical identity remain limited. We developed a genetically encoded probe to identify glutamatergic synaptic vesicles at the levels of both light and electron microscopy (EM) by fusing the mini singlet oxygen generator (miniSOG) probe to an intra-lumenal loop of the vesicular glutamate transporter-2. We then used a 3D imaging method, serial block face scanning EM, combined with a deep learning approach for automatic segmentation of labeled synaptic vesicles to assess the subcellular distribution of transporter-defined vesicles at nanometer scale. These tools represent a new resource for accessing the subcellular structure and molecular machinery of neurotransmission and for transmitter-defined tracing of neuronal connectivity.

## Introduction

The bulk of synaptic neurotransmission is mediated by a surprisingly small group of recycling neurotransmitters, which include the amino acid transmitters, monoamines, purines, and acetylcholine. A defining feature of these neurotransmitters is their ability to package and recycle at the nerve terminal in a process dependent on vesicular neurotransmitter transporters that catalyze synaptic vesicle filling (Edwards, 2007). Because the vesicular transporters are necessary for filling vesicles with specific neurotransmitters, their presence can be used to define synapses by neurotransmitter content.

Synaptic dysfunction is a common feature of neuropsychiatric diseases. For example, a hallmark of age-related neurodegenerative diseases such as Alzheimer’s and Parkinson’s disease is synaptic injury, and aggregation of key proteins that participate in synaptic neurotransmission are thought to be responsible for disease initiation, ultimately leading to cell loss (Burke and O’Malley, 2013; Kanaan et al., 2013). Maladaptive plastic changes in synapse structure and function also underlie key pathological alterations of behavioral and mood disorders ranging from addiction to depression, as well as neurodevelopmental diseases such as schizophrenia or autism spectrum disorders (Kauer and Malenka, 2007; Penzes et al., 2011). Since structure and function of synapses are a key focus across a range of neuroscience disciplines, new tools to study their features within specific neural circuits of interest are sorely needed. Indeed, current tools to assess the structural organization of synapses of defined cell types are not readily compatible with state-of-the-art 3D volume approaches such as serial block-face scanning electron microscopy (SBEM). Additional limitations are imposed by a lack of adequate computational tools for quantitative assessment of these massive datasets. However, advances in molecular genetics, optical imaging, engineering and computing provide new opportunities to develop information-rich strategies to peek into the synapse at high resolution. Here, we combine such advances to achieve a new state-of-the-art platform in imaging and analyzing microcircuit connectivity and synapse structure within neurotransmitter-defined neural networks.

Specifically, we sought to visualize glutamatergic synapses containing the vesicular glutamate transporter-2 (VGLUT2) in projections from the ventral tegmental area (VTA) of the mouse. The VTA is a heterogenous midbrain structure containing dopaminergic, GABAergic and glutamatergic neurons, and VTA dysfunction is associated with numerous neuropsychiatric disorders (Fields et al., 2007). Subpopulations of VTA neurons co-express more than just one neurotransmitter. For example, some VTA neurons express both VGLUT2 and the vesicular monoamine transporter (VMAT2) and release glutamate and dopamine (Chuhma et al., 2004; Hnasko et al., 2010; Kawano et al., 2006; Stuber et al., 2010; Tecuapetla et al., 2010; Yamaguchi et al., 2015); while others express VGLUT2 and the vesicular GABA transporter (VGAT) and release both glutamate and GABA (Root et al., 2014; Yoo et al., 2016). The function of VTA dopamine neurons in reward learning and motivation is well known, but VTA glutamate neurons have also been implicated in processes regulating reward, approach and avoidance behaviors (Alsio et al., 2011; Mingote et al., 2019; Root et al., 2018; Yoo et al., 2016; Zell et al., 2020). Like VTA dopamine neurons, VTA glutamate neurons project to nucleus accumbens (NAc), frontal cortex, and amygdala, but also project to forebrain regions that receive little dopamine input including ventral pallidum (VP) and lateral habenula (LHb) (Hnasko et al., 2012; Taylor et al., 2014). We targeted these VTA glutamate circuits using a conditional viral approach to validate the detection of mini singlet oxygen generator (miniSOG)-labeled VGLUT2-containing synaptic vesicles at nanometer scale by 3D EM combined with deep learning-assisted automated segmentation of labeled and unlabeled synaptic vesicles and other cellular components within pre-synaptic boutons.

## Results

### A viral vector for cell-type specific expression of VGLUT2:miniSOG

In order to identify glutamatergic synapses at nanoscale resolution by 3D EM, we generated a fusion of VGLUT2 and miniSOG, a small fluorescent protein that generates singlet oxygen when activated by intense blue light to locally catalyze the polymerization of diaminobenzidine (DAB) into an osmiophilic reaction product, easily detectable by EM (Shu et al., 2011) (**Figure 1A**). To distinguish labeled from unlabeled vesicles, we designed the new probe to concentrate and accumulate electron dense signal inside synaptic vesicles by inserting miniSOG into the first intralumenal loop of VGLUT2 (**Figure 1A**). We next packaged VGLUT2:miniSOG into a Cre-dependent Adeno-associated virus vector (AAV) and applied it to primary neuronal cultures together with an AAV-Cre:EGFP. Following photooxidation and EM processing, electron dense synaptic vesicles were readily observed, consistent with expected VGLUT2:miniSOG expression and trafficking to synaptic vesicles (**Figure 1B-C**). Negative control neurons transduced with AAV-Cre:EGFP only, did not accumulate osmiophilic DAB precipitates in synaptic vesicles (**Figure 1D**). To confirm the specificity of VGLUT2:miniSOG labeling within the lumen of the vesicles, we used DAB conjugated with lanthanide chelates (cerium) (Adams et al., 2016). This allowed us to uniquely differentiate the spectral signal of cerium in the DAB deposits (**Figure 1E-F**) by imaging electron energy-loss spectroscopy (EELS) in an energy-filtered transmission EM, providing confidence that the DAB signal could be clearly resolved with the VGLUT2:miniSOG-labeled vesicles.

**Figure 1.**
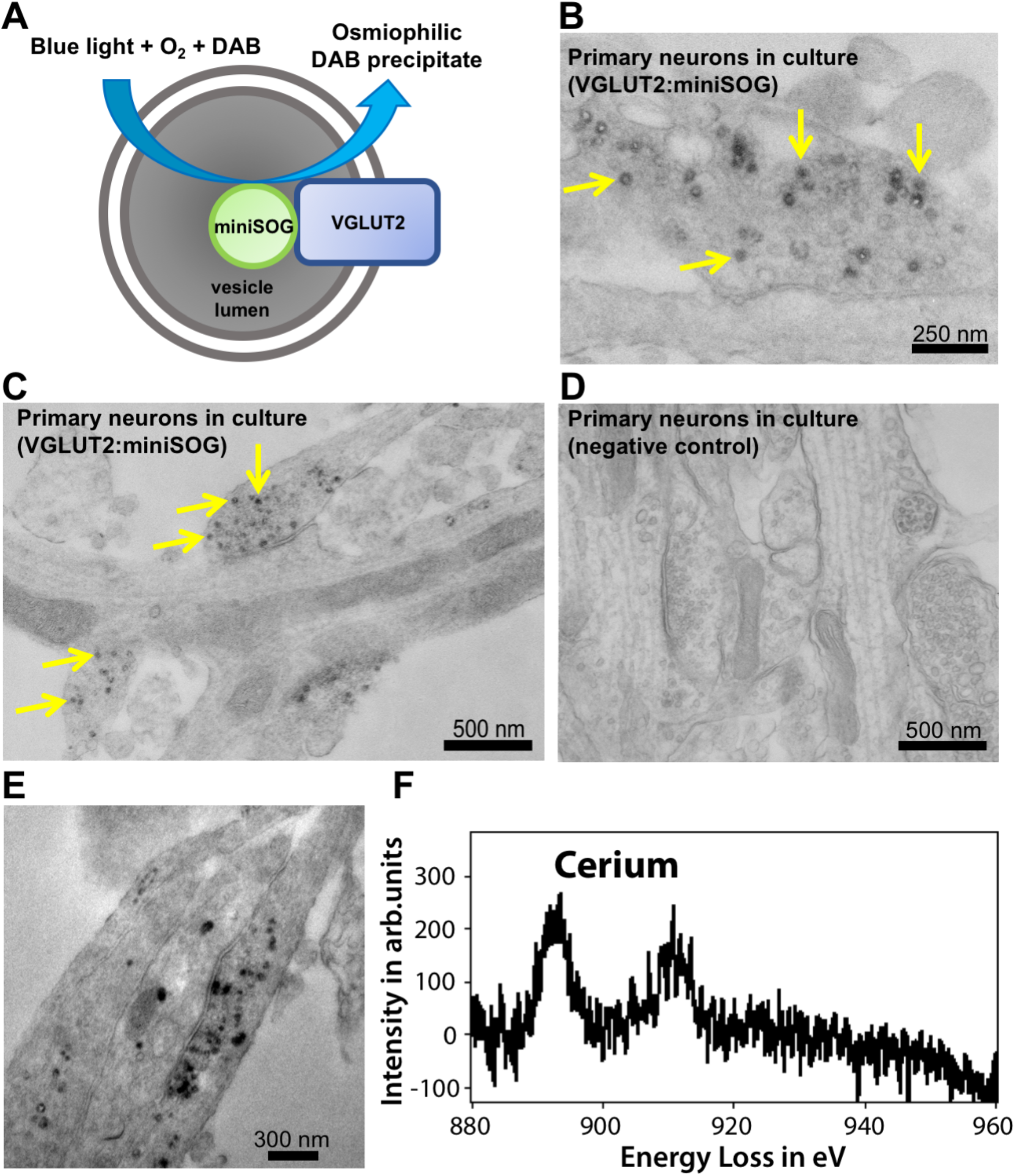
Expression and visualization of VGLUT2:miniSOG-labeled vesicles in primary neuronal cultures. **(A)** Schematic of VGLUT2:miniSOG in a synaptic vesicle; intralumenal location of miniSOG which in the presence of blue light and oxygen induces polymerization of DAB into an osmiophilic precipitate within the vesicle lumen.-**(B, C)** Electron micrographs showing VGLUT2:miniSOG-labeled vesicles (yellow arrows) in cultured primary neurons transduced with AAV-DIO-VGLUT2:miniSOG plus AAV-Cre:eGFP or **(D)** AAV-Cre:eGFP only (negative control). (**E**) TEM image of primary rat cortical neuron expressing VGLUT2:miniSOG. The darker signal reflects the VGLUT2-specific DAB labeling within the vesicles and (**F**) corresponding EELS spectrum showing the Cerium signal.

### VGLUT2:miniSOG rescues glutamate release from neurons lacking VGLUT2

In order to test whether VGLUT2:miniSOG is useful for long-range ultrastructure circuit mapping while retaining its function, we injected AAV-DIO-VGLUT2:miniSOG together with a conditional viral vector to express Channelrhodopsin-2 (AAV-DIO-ChR2:mCherry) into the VTA of DAT-Cre conditional knockout (cKO) mice lacking endogenous VGLUT2 (*Slc6a3^+/Cre^; Slc17a6^flox/flox^*) (**Figure 2A-B**). After 4-6 weeks we performed *ex vivo* whole-cell recordings from acute brain slices containing the nucleus accumbens (NAc) medial shell and used 5-ms pulses of blue light to activate ChR2 and evoke neurotransmitter release from dopamine axons. Consistent with prior results (Bimpisidis et al., 2019; Steinkellner et al., 2018; Stuber et al., 2010; Wang et al., 2017), we measured no optogenetic-evoked glutamate EPSCs in cKOs; however, cKO mice with Cre-dependent expression of VGLUT2:miniSOG in dopamine neurons showed robust AMPA- and NMDA-driven opto-triggered EPSCs (**Figure 2C**). Furthermore, immunostaining for miniSOG indicated that VGLUT2:miniSOG could be detected in Tyrosine hydroxylase (TH)-expressing VTA dopamine neurons as well as their synaptic terminals in medial NAc shell (**Figure 2D**).

**Figure 2.**
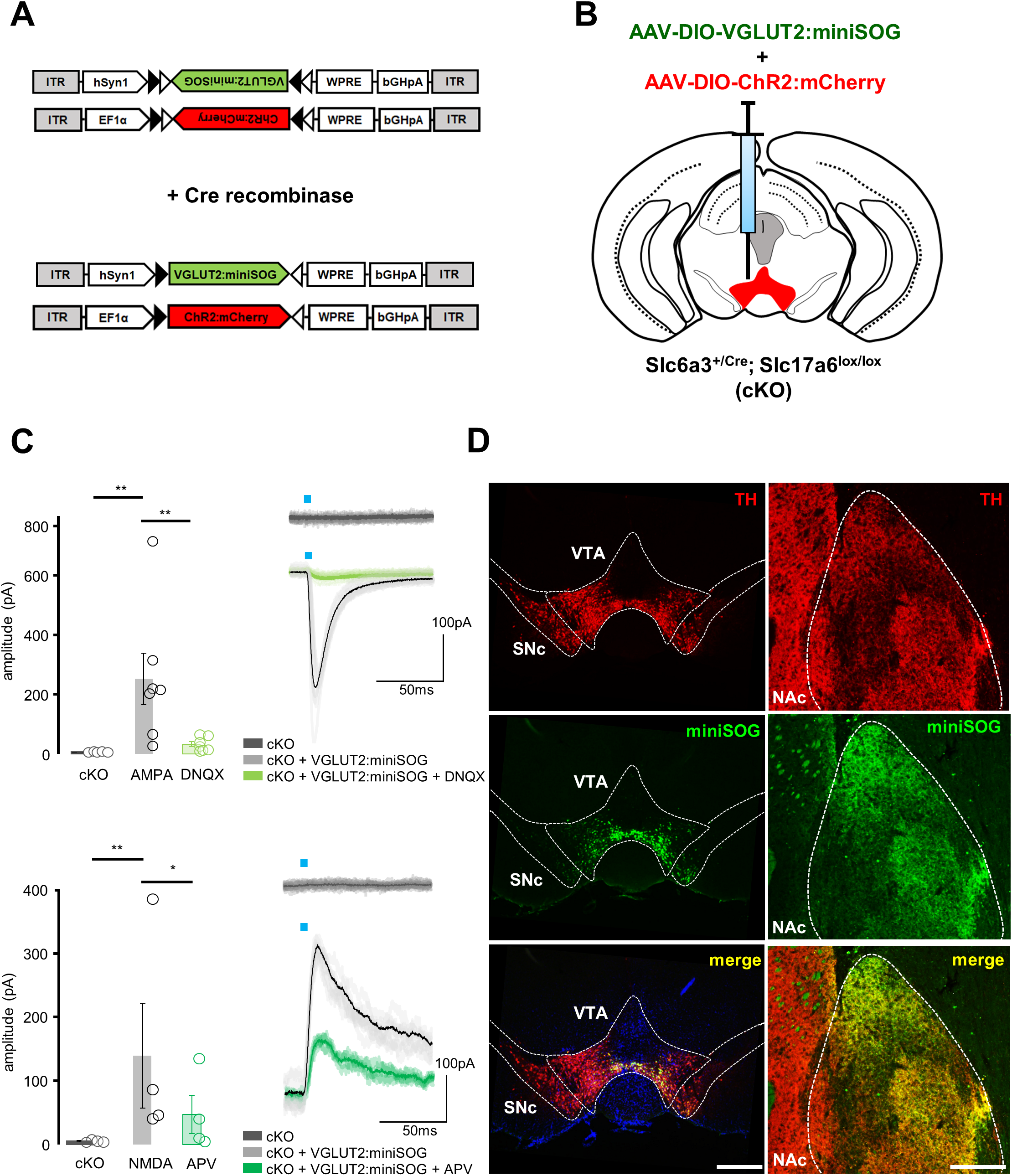
VGLUT2:miniSOG rescue of glutamate co-release in mice lacking endogenous VGLUT2 in dopamine neurons. **(A)** Dual injection of Cre-dependent AAV to express VGLUT:miniSOG and ChR2:mCherry (**B**) into the VTA of cKO mice lacking endogenous VGLUT2 in dopamine neurons. **(C)** Expression of VGLUT2:miniSOG in VTA dopamine neurons rescued ChR2-evoked postsynaptic AMPA-(top) or NMDA-receptor (bottom) mediated EPSCs recorded from postsynaptic neurons in NAc medial shell; scale 100 pA, 50ms; *p<0.05, **p<0.01 by paired t-test. **(D)** VGLUT2:miniSOG expression was detected in TH-expressing dopamine neurons using an anti-miniSOG antibody, both at the injection site (VTA) and in medial NAc shell terminals. Coronal sections, scale 500μm (VTA), 200μm (NAc).

We next used a similar approach to express VGLUT2:miniSOG in VTA glutamate neurons using VGLUT2-Cre mice. Three weeks after AAV injection we used immunohistochemistry against miniSOG and detected punctate expression of miniSOG-labeled VGLUT2 in major projection areas of these cells including the medial NAc shell, VP, and LHb (Hnasko et al., 2012; Taylor et al., 2014) (**Figure 3A**). We confirmed that an antibody directed against endogenous VGLUT2 co-localized with miniSOG, and that VGLUT2:miniSOG localized with the vesicular marker synaptophysin (**Figure 3B**), consistent with the trafficking of VGLUT2:miniSOG to synaptic vesicles.

**Figure 3.**
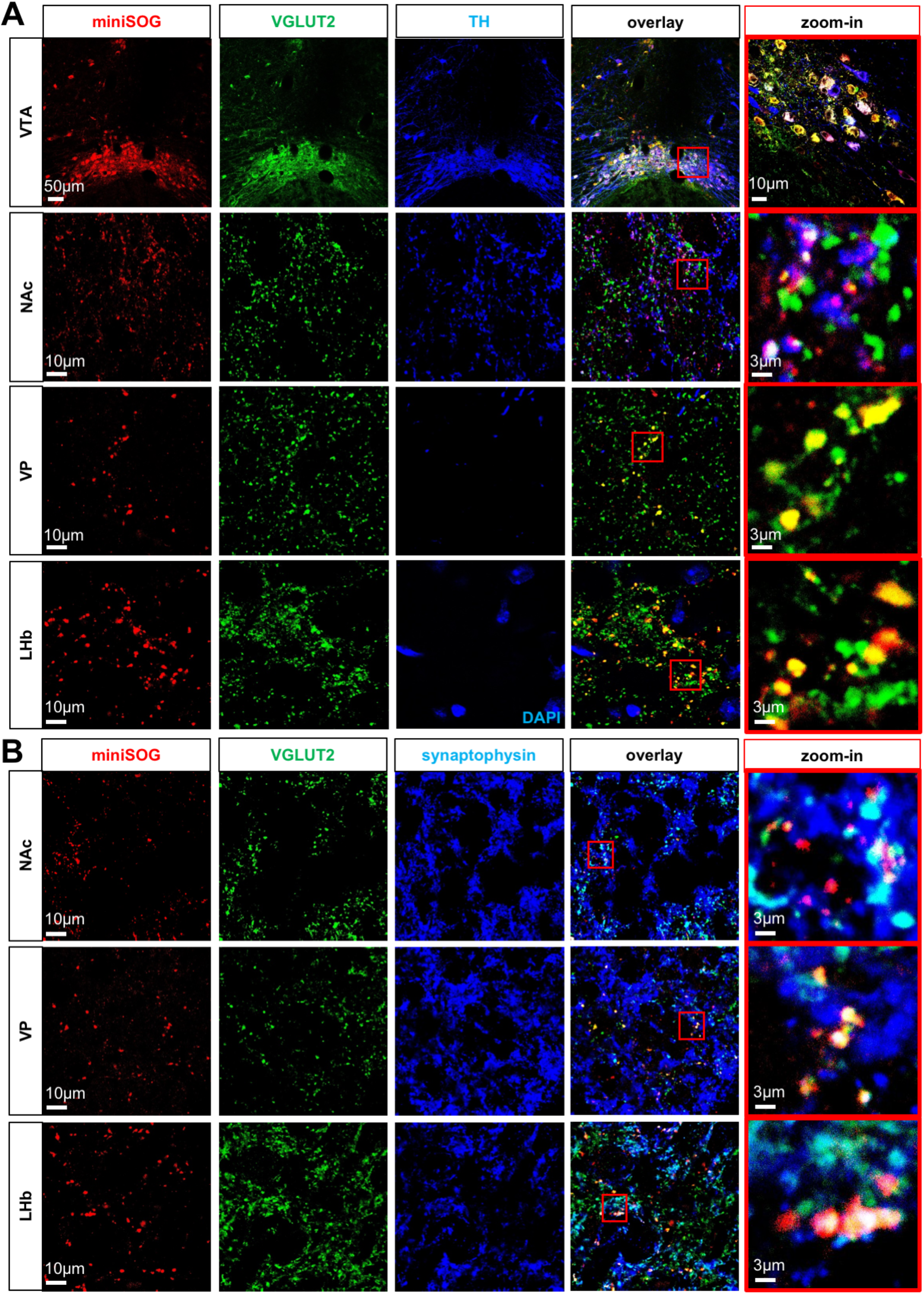
VGLUT2:miniSOG traffics and localizes to synaptic terminals of VTA glutamate neurons. VGLUT2-Cre mice were injected with AAV-DIO-VGLUT2:miniSOG into the VTA and confocal images were made through coronal sections in VTA, NAc, VP and LHb. **(A)** Immunostaining against miniSOG and VGLUT2 demonstrates co-localization of VGLUT2 and miniSOG as well as partial co-localization with the dopamine marker TH in a subpopulation of the VTA projections to NAc, but not VP. DAPI (blue) is shown in LHb due to lack of TH signal here. **(B)** miniSOG and VGLUT2 also co-localize with the synaptic vesicle marker synaptophysin. Images with red frame in the right column are higher magnification.

### Detection of VGLUT2:miniSOG in synaptic vesicles in mouse brain

We next tested for VGLUT2:miniSOG labeling at presynaptic vesicles at the ultrastructural level by transmission electron microscopy (TEM). Three weeks after injection of AAV-DIO-VGLUT2:miniSOG, mice were perfused with fixative, brains were extracted and slices through NAc were exposed to intense blue light illumination in the presence of DAB before processing samples for conventional TEM. Electron micrographs from boutons showed intense DAB labeling in the lumen of presynaptic vesicles (**Figure 4A**). Most importantly, because the DAB reaction product was contained within vesicles, it allowed for easy identification of labeled presynaptic boutons without masking other subcellular details, such as the active zone or postsynaptic densities.

**Figure 4.**
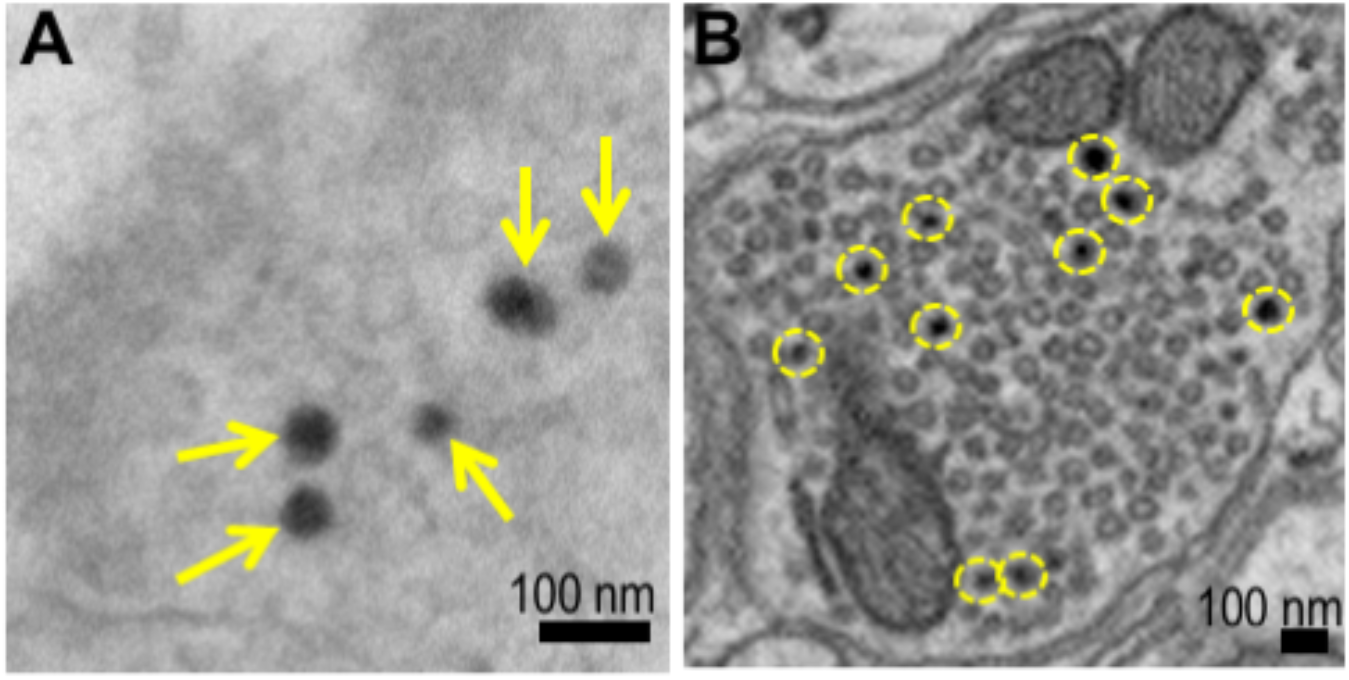
EM visualization of VGLUT2:miniSOG-positive vesicles in brain. **(A)** Electron micrograph of a presynaptic bouton from the NAc expressing VGLUT2:miniSOG showing the intense DAB labeling in the lumen of presynaptic vesicles (yellow arrows). The sample was processed by conventional TEM with minimal staining (only osmium tetra-oxide) so that the specific VGLUT2:miniSOG EM signal would be generated only by passage in osmium to add electron density to the DAB precipitates. **(B)** Analysis using SBEM in NAc revealed DAB-labeled osmiophilic reaction product fills synaptic vesicles after photooxidation of VGLUT2:miniSOG (yellow circles).

In order to image the 3D morphology of these glutamate-defined synapses we utilized SBEM. This approach allows imaging the pattern of release sites at both the neural circuit and sub-cellular (synaptic) level. The specimen preparation involves miniSOG-mediated DAB oxidation, subsequent processing with heavy metal staining and imaging by scanning electron microscopy (Wilke et al., 2013). Similar to our results with conventional TEM, we observed synapses labeled by dark, defined electron-dense vesicles that could be easily distinguished from neighboring unlabeled vesicles (**Figure 4B**). Of note, not all vesicles in a presynaptic bouton contained DAB reaction product.

### Analysis of VTA glutamate synapses by projection target

The use of large-scale SBEM data requires accurate segmentations based on grey level images and which respect the complexity and multi-scale nature of these data. However, identifying and extracting objects from within these large and complex 3D image volumes, especially synaptic vesicles which can number in the thousands per bouton, can be a challenging task. We thus applied a recently established pipeline for EM auto-segmentation and analysis, CDeep3M (Haberl et al., 2018) and implemented a workflow for image processing and analysis to allow rapid and accurate quantification of pre-synaptic ultrastructural details and 3D synaptic morphology (**Figure 5**). CDeep3M uses a 3D convolution neural network architecture which takes an image as an input (**Figure 5A**), and passes that data through a series of convolutional layers that discriminate patterns in the grey level pixel features of an object of interest, and returns an output that represents the probability of input pixels belonging to that certain object. In order for the neural network to discriminate a specific object, ground-truth labeled data is provided and the network is trained to recognize the input by weighting the connections between convolutional layers. A trained network can then be scaled to predict the presence of objects in the large fields of SBEM image data (**Figure 5B**). On these predictions we can perform morphometric data analysis that allows rigorous quantitative assessment of differences across brain areas. As proof-of-principle we imaged three different major release sites of VTA glutamate projection neurons within the same VGLUT2-Cre mouse expressing VGLUT2:miniSOG in VTA neurons – the medial NAc shell, VP and LHb. Separate sections through these regions were photo-oxidized and processed for SBEM. Images were acquired with a z-step of 24 or 30 nm, and pixel size of 2.4-2.5 nm, allowing us to screen volumes 3,230 to 9,700 μm^3^ with high spatial resolution. VGLUT2:miniSOG-labeled vesicles were detected within each of these three projection targets of VTA glutamate neurons. We identified and counted miniSOG-labeled vesicles, as well as unlabeled vesicles in boutons containing at least one labeled vesicle, for all three brain areas examined (**Figure 5C**, **Figure 6A-C**). Approximately 40% of the synaptic vesicles in VP boutons were VGLUT2:miniSOG labeled, compared to ~10% in LHb and NAc (**Figure 5C**).

**Figure 5.**
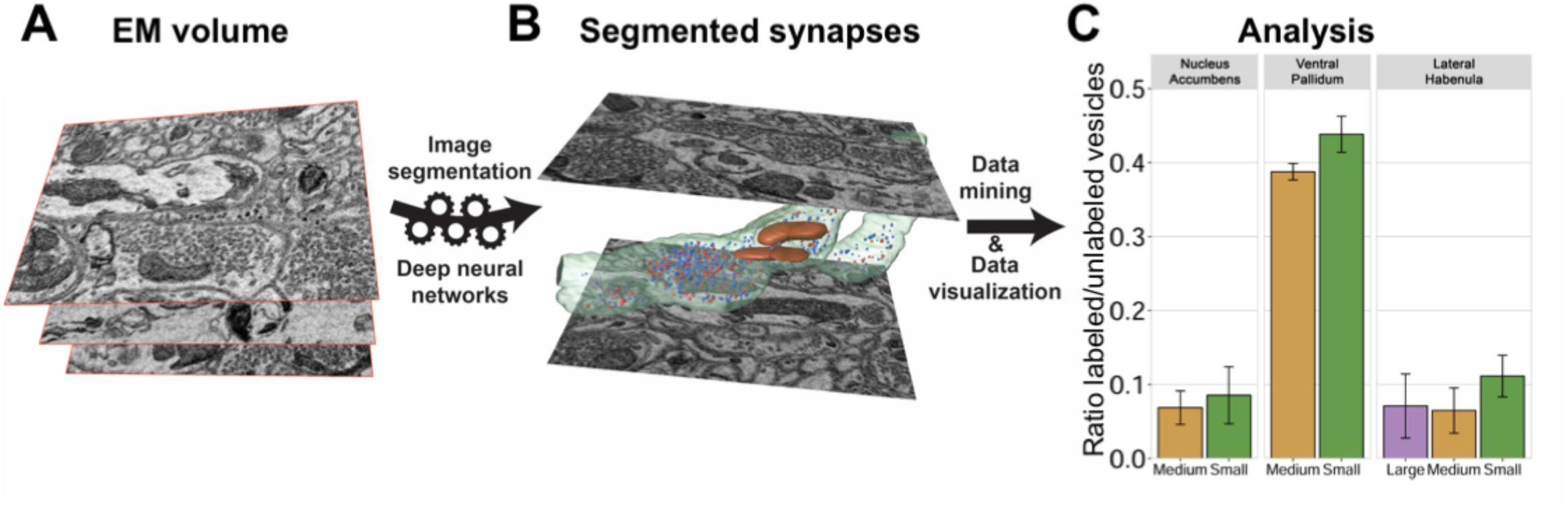
Workflow for image processing and analysis for quantification of pre-synaptic ultrastructural detail. CDeep3M allows large-scale automatic image segmentation using state-of-the-art deep neural networks. **(A)** The process of automated labeling of an image volume requires training the network using a subvolume containing ground-truth segmentations of the object-of-interest. The trained model is then used to predict the occurrence of the object across the whole image volume. **(B)** The model’s prediction can then be 3D-reconstructed into image objects, such as VGLUT2:miniSOG-labeled vesicles (red), unlabeled vesicles (blue), and mitochondria (orange). Axons and their constituent boutons were traced by human annotators and rendered translucently (green). **(C)** A custom post-processing pipeline was used to identity individual objects and extract metrics of interest. Ratio of VGLUT2:miniSOG-labeled synaptic vesicles to the total amount of vesicles inside a presynaptic bouton in three different VTA-projection areas. Bouton sizes: large ≥ 1 μm^3^; medium 0.5 to 1 μm^3^; small < 0.5 μm^3^.

**Figure 6.**
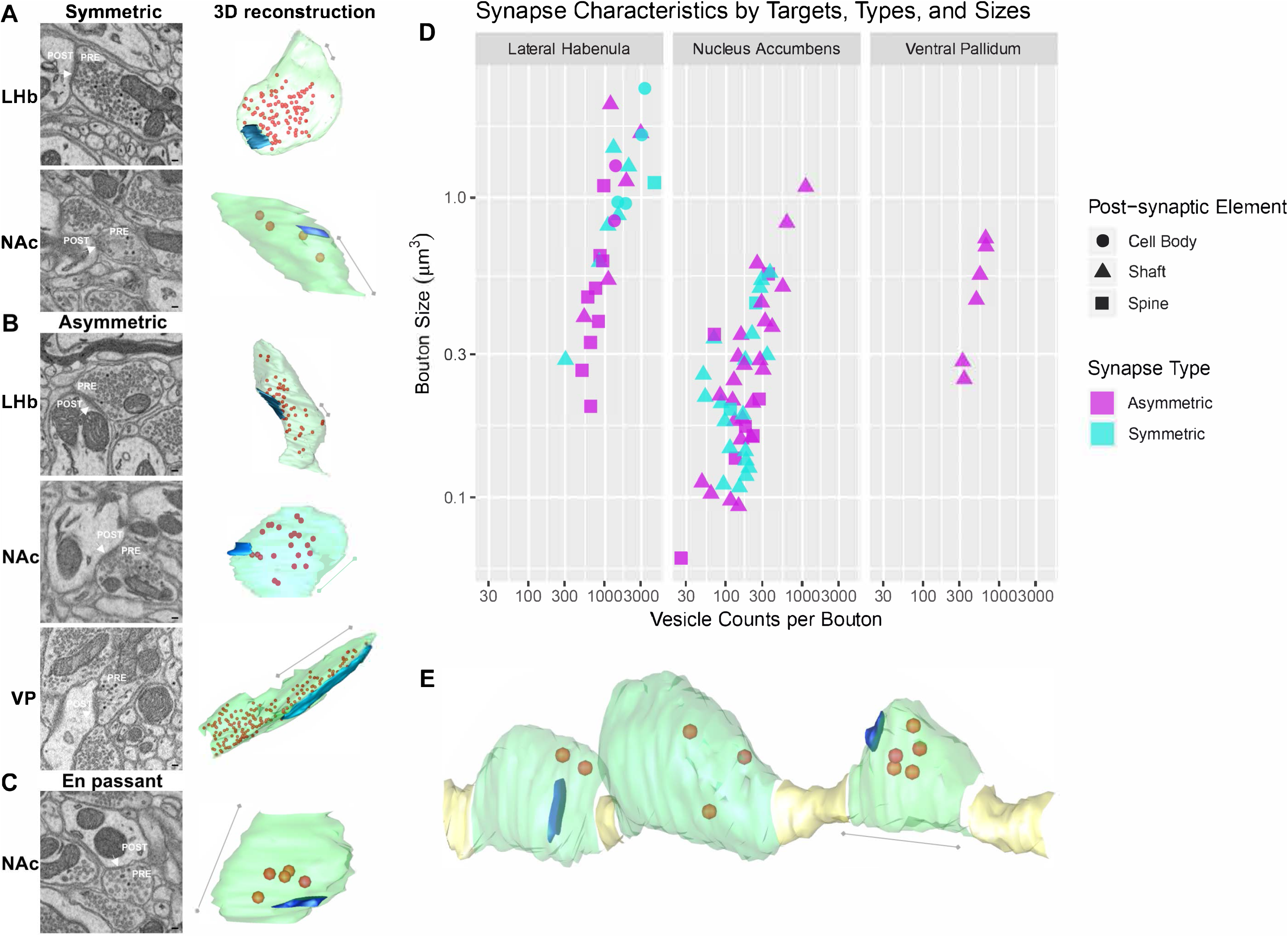
Synapse characteristics by target, type, and size. **(A-C)** Electron micrographs showing different types of VGLUT2:miniSOG-positive boutons (labeled PRE) in three brain areas and their corresponding 3D reconstruction showing the plasma membrane in green, VGLUT2:miniSOG-labeled vesicles in orange, and active zone in blue. Synapses (white arrows) were visually indexed as to whether they were asymmetric or symmetric and if the post-synaptic target (labeled POST in the micrographs) was a cell body, dendritic shaft, or dendritic spine via human annotation. Asymmetric synapses have pronounced post-synaptic densities while symmetric synapses contained relatively equal density gradients at pre and post synaptic side. Scale bars, 100 nm for EM micrographs and 500 nm for 3D reconstructions. **(D)** Bouton sizes measured by summing the volume inside the bouton’s plasma membrane are plotted against vesicle counts per bouton, including both unlabeled and VGLUT2:miniSOG-labeled, in each of the three areas analyzed. Boutons are distinguished by post-synaptic target type and synapse. Boutons which contained synapses that were either indistinguishable, orientated *en face* to the cut plane, contained multiple release sites on different targets, or contained multiple release sites that contained asymmetric or symmetric properties were omitted. **(E)** *En passant* varicosities are abundantly observed in the nucleus accumbens. Scale bar, 500 nm.

As shown in **Figure 6**, the majority of VGLUT2:miniSOG-positive boutons from VTA on to LHb were on dendrites (**Figure 6D**). This included dendritic shafts (11 of 27 complete, labeled boutons) and 5 of these were asymmetric while 6 were symmetric, as well as dendritic spines (10 of 27) where the majority were asymmetrical (9 of 10). A smaller proportion were on neuronal cell bodies (6 of 27) where 4 of 6 were symmetrical. Though we detected fewer overall boutons in the VP from this animal, all the VGLUT2:miniSOG-positive boutons identified were asymmetric and on dendritic shafts (6 of 6). In contrast, most of the vesicles in the NAc were identified along axons in which the *en passant* varicosities transversed the image volume without terminating (**Figure 6E**). These varicosities frequently did not associate with a discernable post-synaptic compartment (55 of 107). In some of these cases the varicosities were surrounded exclusively by other axons, in other cases the sectioning plane was parallel to the plane of the synaptic contact (en face) making classification impractical. However, for those VGLUT2:miniSOG-containing varicosities or boutons with adjacent post-synaptic compartments, the majority were on dendritic shafts (43 of 107) with fewer on dendritic spines (9 of 107).

Given the high quality ultrastructural preservation and the DAB labeling clearly confined to the lumen of the VGLUT2:miniSOG-labeled vesicles, subcellular details were easily visualized allowing for analysis of the synaptic vesicles and intracellular organelle profiles in labeled presynaptic boutons for each of the three projection areas (**Figure 7**). The lateral habenula presented the biggest boutons (average bouton sizes: 0.87 μm^3^ in LHb, 0.24 μm^3^ in NAc, 0.43 μm^3^ in VP, **Figure 7A**). We also measured the diameter of labeled and unlabeled vesicles in each region, which revealed that labeled and unlabeled vesicles differed little in size (**Figure 7B**). We then calculated the total volume of each bouton, the total mitochondrial volume present in each bouton, and aggregate volume of labeled and unlabeled synaptic vesicles (**Figure 7C**). A similar fraction of volume/bouton in all three brain areas was occupied by mitochondria (15.8% volume fraction in LHb, 16.5% in NAc, 10.4% in VP) and we identified at least one mitochondrion in more than 75% of boutons in each terminal region: LHb (23/27), NAc (81/107), and VP (5/6). Volume fraction occupied by synaptic vesicles was a mean of 4% in LHb, 2.2% in NAc, and 2.5% in VP.

**Figure 7.**
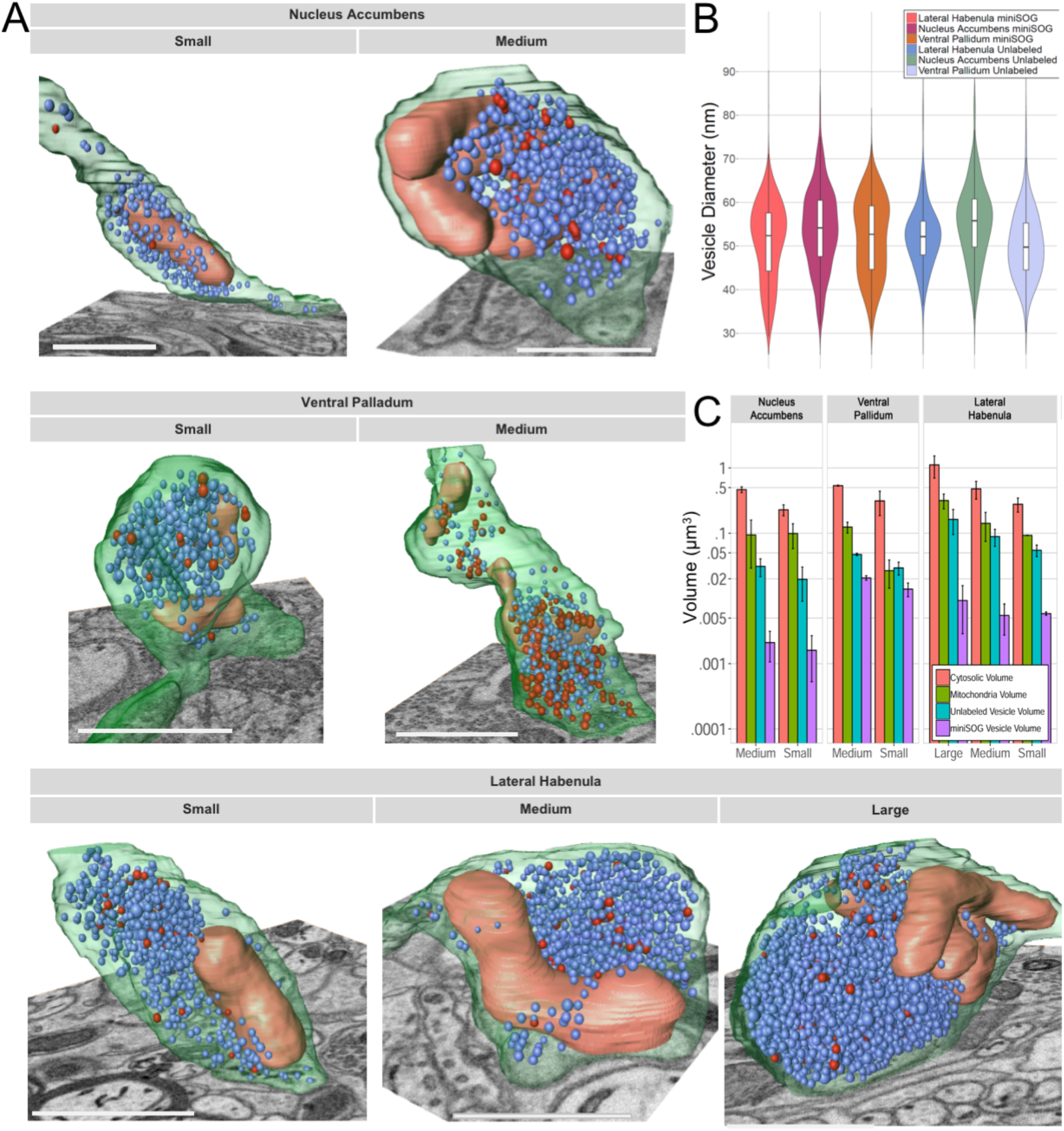
Profiles of synaptic vesicles and boutons across projection targets. (**A**) 3D reconstructions of presynaptic boutons from three different VTA projection areas show the distribution of VGLUT2:miniSOG-labeled (red) and unlabeled synaptic vesicles (blue), plasma membrane (green) and mitochondria (orange). Scale bars, 1 micron. (**B**) The diameter of VGLUT2:miniSOG-labeled and unlabeled vesicles was determined for each of the three brain areas examined as shown in the violin plot, in which the standard box plot displays the mean, range, and interquartile range while being enveloped by a mirrored and colored relative frequency histogram. **(C)** Profile of presynaptic organelles in different brain regions. Total volumes of organelles in presynaptic boutons: total vesicle volume was measured by summing the spherical volumes of every vesicle per bouton; cytoplasmic volume was determined by subtracting the mitochondria and vesicle volumes from the volume inside the bouton membrane.

We further analyzed the distribution of synaptic vesicles of different sizes in the three different brain areas relative to their distance from the active zone. The vast majority of vesicles were localized within 1 micron of the active zone (89.5% of all vesicles, 84.1% for VGLUT2:miniSOG-labeled and 89.9% for unlabeled vesicles). VGLUT2:miniSOG-labeled vesicles do not appear to be preferentially clustered in proximity to an active zone but rather scattered within the larger pool of unlabeled vesicles (**Figure 8A-B).** In LHb and NAc, a subset of vesicles can be observed in an outlier cluster at approximately 80 nm in vesicle diameter (**Figure 8B**). Interestingly, vesicles in the NAc and LHb showed two major clusters localized within 1 micron and more distant subpopulations localized ~1.5 and 2 microns from the active zone, respectively, suggesting these might represent different vesicle pools. Overall, for all three areas examined, no apparent correlation was observed between the size of the vesicles and their distance from the active release site.

**Figure 8.**
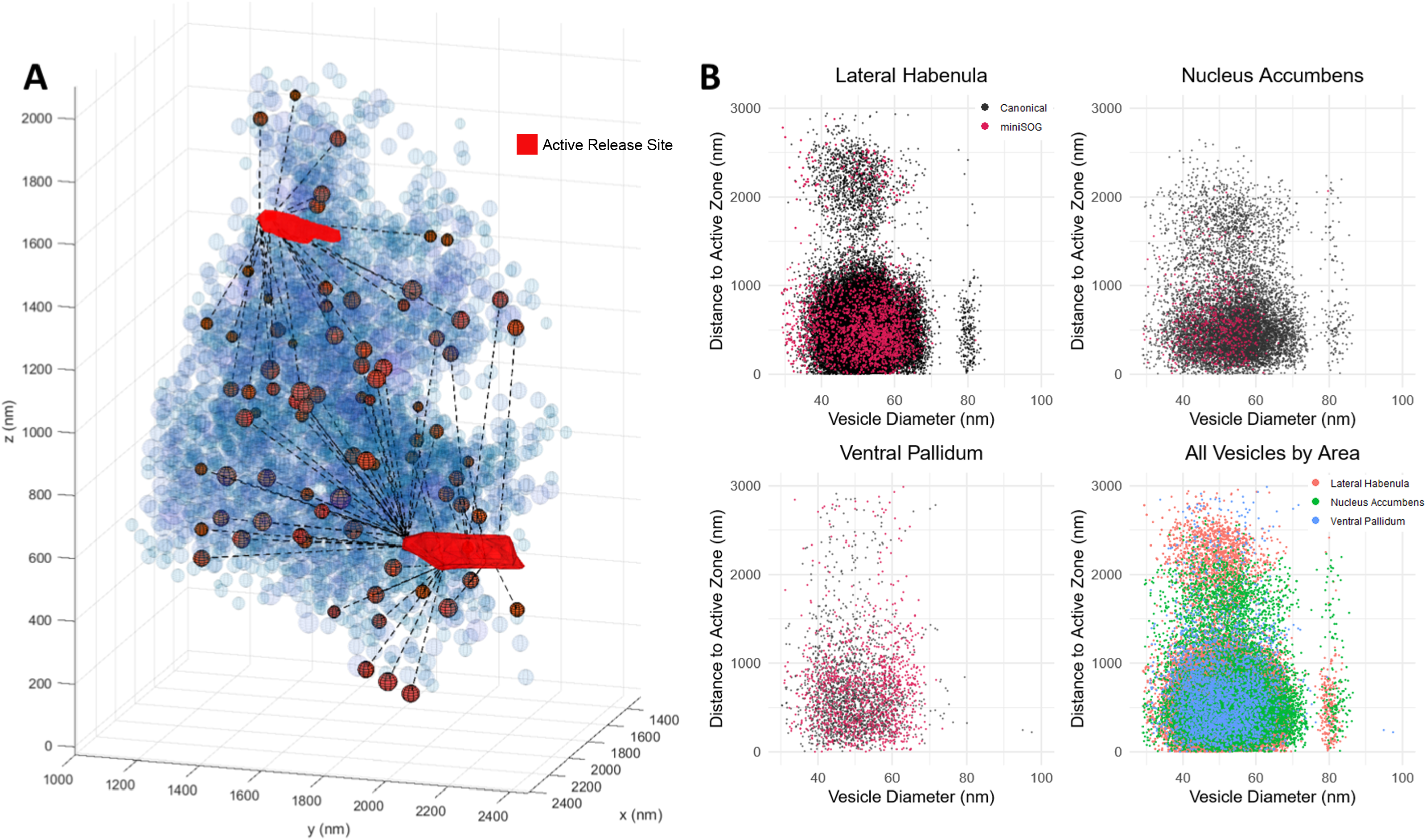
Spatial distribution of synaptic vesicles relative to the active zone. (**A**) Distance to active zone visualization of a presynaptic bouton from LHb showing unlabeled presynaptic vesicles (faded in blue) and VGLUT2:miniSOG-labeled vesicles (in orange) plotted with line representing the shortest distance to the nearest active zone, utilizing a k-nearest neighbors algorithm. The active zone was identified in the EM micrograph by docking of vesicles to the plasma membrane (red). (**B**) Distribution of VGLUT2:miniSOG-labeled (in red) and unlabeled (canonical, in black) synaptic vesicles of different sizes in the different brain areas with respect to their 3D distance from the active zone.

## Discussion

By introducing the miniSOG EM tag *in vivo* we were able to assess the spatial distribution of glutamatergic synapses, and vesicle pools in circuits established by VTA glutamate neurons at nanometer resolution by 3D EM. Genetically encoded EM-compatible tags such as miniSOG can be directly fused to a protein of interest and all reagents needed to render them EM visible are small molecules that readily permeate well-fixed tissue, therefore there is no conflict between detectability of the protein target and excellent ultrastructural preservation of cellular landmarks. Our approach allows for high resolution 3D reconstruction circumventing limitations encountered with more conventional methods such as antibody-based techniques, which to date represent the most common methods for imaging molecular-defined synapses. Even though immuno-detection of proteins in fixed tissue continues to support innumerable advances, the approaches have severe drawbacks, particularly at the ultrastructural level. Conventional immuno-electron microscopy (EM) techniques have fundamental unresolved problems with the limited ability of most antibodies to recognize heavily fixed antigens and to penetrate into and wash out of well-fixed cells and tissues. The fixation, embedding and sectioning required for these techniques can reduce availability of most of the epitopes resulting in low sampling rates. Additionally, immuno-EM requires use of detergents to gain access to intracellular epitopes and this creates a challenge in keeping the ultrastructure intact while retaining antigenicity for a given antibody.

Various super-resolution optical methods have been developed to improve spatial resolution towards that of EM while exploiting the relative ease with which target proteins can be specifically tagged with fluorophores. However, super-resolution fluorescence only allows to visualize the specifically tagged proteins against a featureless black background, requiring numerous immune-labeling combinations to reconstitute a larger context of the cellular environment. The introduction of genetically targetable EM tags such as miniSOG (Shu et al., 2011), APEX2 (Lam et al., 2015), or HRP (Atasoy et al., 2014) circumvents these issues allowing 3D visualization of proteins of interest around the complex subcellular, cellular, and tissue morphology under investigation. We expect that our new genetic labeling strategy and subsequent analysis techniques will enable new research directions looking at neuronal neurotransmitter profiles, studying the composition and structural organization of synaptic boutons.

As proof-of-principle, we tagged the vesicular neurotransmitter transporter VGLUT2 with miniSOG for several reasons. First, tagging VGLUT2 at an intralumenal loop allows for accumulation of DAB precipitates inside vesicles and hence eases signal discrimination against EM background. Second, because label is restricted to vesicles, surrounding ultrastructure is preserved and easy to distinguish. Third, the presence of VGLUT2 provides additional information, specifying the synapse as capable of glutamatergic transmission. Indeed, VGLUT2 localizes primarily to synaptic vesicles present in presynaptic terminals which makes it an ideal protein for visualizing glutamatergic synapses (Fremeau et al., 2001). Further, VGLUT2 is both, necessary and sufficient to enable synaptic vesicles to store and release glutamate and is the main vesicular glutamate transporter expressed by non-telencephalic neurons in the mammalian brain (Fremeau et al., 2001; Herzog et al., 2001; Takamori et al., 2001).

To demonstrate the capacity of VGLUT2:miniSOG to label synaptic vesicles *in vivo* we targeted it to VGLUT2-Cre-experessing VTA neurons. The VTA is a midbrain structure well known for its dopamine neurons and role in the regulation of motivated behaviors. While a subset of VTA DA neurons also express VGLUT2 enabling the co-release of glutamate, a larger population of VTA glutamate neurons lack dopamine neuron markers, also contribute to reward processes, but have been minimally characterized in terms of synapse architecture. We demonstrate that VGLUT2:miniSOG targeted to the VTA colocalized with VGLUT2 and synaptophysin in efferent targets including NAc, LHb, and VP indicating that VGLUT2:miniSOG traffics to presynaptic terminals, where we also demonstrate it can rescue optogenetic-evoked glutamate release. Photooxidation and subsequent EM through medial NAc shell, VP, and LHb resulted in concentration of DAB precipitate within distinct, membrane-enclosed subcellular structures resembling synaptic vesicles and preserved synaptic ultrastructure. This indicates that VGLUT2:miniSOG trafficked to synaptic vesicles of boutons in the investigated projection areas. Of note, we also detected fluorescent signal in both VTA glutamate and dopamine neuron cell bodies. Whether this represents presence of the transporter processing through the secretory pathway, or mis-localization and accumulation is not known, but represents an important potential caveat to the AAV driven over-expression approach.

Despite the overexpression of VGLUT2:miniSOG it is remarkable that only a minority of apparent synaptic vesicles in identified presynaptic structures contained DAB label, even with the high permeability of the needed reaction factors DAB, oxygen and light. Because sup-populations of VTA glutamate neurons co-release dopamine or GABA, some of these vesicles may instead contain the type-2 vesicular monoamine transporter (VMAT2) or the vesicular GABA transporter (VGAT) (Hnasko et al., 2010; Kawano et al., 2006; Root et al., 2014; Yoo et al., 2016). However, it may also be the case that many or most of the unlabeled synaptic vesicles are devoid of any vesicular transporter, though the biological consequences of such a configuration is not clear. Consistent with this, immuno-EM studies examining VGLUT2 in VTA terminals showed only a small fraction of synaptic vesicles labeled with VGLUT2 (Zhang et al., 2015) within pre-synaptic compartments, though here it may be related to inherent trade-offs associated with immuno-EM that resulted in partial epitope detection (Root et al., 2018; Root et al., 2014).

As noted, a subset of VTA glutamate neurons are capable of releasing other neurotransmitters depending on projection target. For example, a minority (~10-20%) of all VTA glutamate neurons expresses the dopamine marker TH, however those VTA glutamate neurons that project to the NAc appear to express VGLUT2 at much higher rates (>70%) (Yamaguchi et al., 2011; Zell et al., 2020). Thus most of the synapses we assessed in NAc are likely to be derived from VTA dopamine neurons that co-express VGLUT2:miniSOG, and this presumably accounts for our widespread identification of *en passant* synapses typical of dopamine neurons in NAc (Descarries et al., 2008; Descarries and Mechawar, 2000; Descarries et al., 1996; Nirenberg et al., 1996; Sesack et al., 1998). However, recent studies have demonstrated that not all of these synapses are competent for vesicular release (Ducrot et al., 2020; Liu et al., 2018), which provides another possible explanation for why only a minority of synaptic vesicles label with VGLUT2 at many of these putative release sites. On the other hand, subsets of VTA glutamate neurons that project to VP and LHb express VGAT and co-release GABA (Yoo et al., 2016). Indeed, this includes ~70% of the VTA glutamate neurons projecting to LHb and these neurons were shown to make both asymmetric and symmetric contacts (Root et al., 2014), a remarkable finding validated by the observations presented herein.

While we focus here on describing VTA glutamate neuron projections that have some specific properties, some features are nonetheless consistent with prior descriptions of other glutamatergic synapses. Presynaptic boutons of excitatory synapses have been depicted to contain round, clear vesicles of diverse size, a general phenomenon described in several species and brain regions (Harris et al., 2015; Qu et al., 2009; Tao et al., 2018). The smaller ones range between ~30 to 50 nm, while other synapses contain also bigger vesicles (>60nm) consisting of clear vesicles as well as dense core vesicles (Harris and Weinberg, 2012). These different vesicles sizes were also observed in our VTA glutamate neurons where the VGLUT2:miniSOG-labeled vesicles range between 30 to 60 nm in diameter in the NAc, and 30 to 70 nm in the LHb and VP. The bigger cluster around 80 nm in diameter was composed of unlabeled vesicles and mostly evident in the LHb and NAc. While most glutamatergic excitatory synapses are formed on dendritic spines with some exceptions, in contrast to GABAergic inhibitory synapses that are primarily formed on dendritic shafts (Tao et al., 2018), we observed the majority of synapses in NAc and VP on dendritic shafts while LHb was evenly distributed between dendritic shafts and spines. Also, the total number of vesicles per bouton (~300-3,500) observed in the LHb exceeded the average number of vesicles described for other brain areas (mean values ~150-500) (Kasthuri et al., 2015; Sando et al., 2017). Interestingly, we observed at least one mitochondrion in more than 75% of boutons in each terminal region, in contrast with what has been described in other brain areas such as in hippocampal CA1 and CA3, where mitochondria occurred in less than 50% of synaptic boutons (Shepherd and Harris, 1998).

To summarize, we here describe a new genetically encodable probe, VGLUT2:miniSOG, which can be specifically targeted to Cre-expressing cells as shown here for either VTA glutamate or dopamine neurons. Our approach enables identification of glutamatergic vesicles using both light and electron microscopy and provides a general pipeline for automated segmentation of labeled synaptic vesicles after 3D EM imaging by SBEM using a recently developed deep learning approach (Haberl et al., 2018). Combining these 3D datasets and the computational analysis paves the way for future studies quantifying the number and distribution of vesicles, together with their neurotransmitter content. This will allow for relating the projection-specific identification of neurotransmitter content across brain areas with diverse synapse organization.

## Materials and Methods

### Mice

Mice were used in accordance with protocols approved by the University of California, San Diego Institutional Animal Care and Use Committee (protocol # S12080). Mice expressing Cre under the control of DAT (*Slc6a3^IRESCre^*, Jackson Stock # 006660) or VGLUT2 (*Slc17a6^IRESCre^*, Jackson Stock # 016963) regulatory elements were obtained from The Jackson Laboratory and then bred in house. Mice lacking VGLUT2 in dopamine neurons (VGLUT2 cKOs) (*Slc17a6^flox/flox^; Slc6a3^+/IRESCre^*) were generated as described previously (Steinkellner et al., 2018). Mice were (>10 generations) backcrossed to C57BL/6 background. Both sexes were used for experiments. Mice were group-housed on a 12-h light-dark cycle, and with food and water available *ad libitum*.

### DNA constructs and adeno-associated viral vectors

Rat *Slc17a6* (VGLUT2) cDNA was a gift from Robert Edwards (UCSF) and pAAV-hSyn1-DIO-VGLUT2:mSOG was cloned using standard molecular biology techniques. Briefly, miniSOG was inserted between glycines 107 and 108 (Foss et al., 2013) flanked by 5’ and 3’ linkers (5’: AGCACGAGCGGGGGATCTGGGGGCACTGGTGGTTCA [amino acids: STSGGSGGTGGS]; 3’: GGAGGTACCGGGGGCACTGGCGGGAGCGGTGGGACCGGC [amino acids: GGTGGTGGSGGTG]) in pUC57 before excision and subcloning of VGLUT2:miniSOG into a pAAV-hSyn1-DIO plasmid using restriction enzymes. All sequences were verified by site sequencing. Plasmid DNA was purified from *E. coli* and DNA was packaged into AAVDJ serotype by the Salk GT3 vector core and titer estimated by qPCR (final titer: 2.27 × 10^13 genome copies per milliliter [gc/ml]) (La Jolla, CA). The plasmid construct was tested for Cre -dependent recombination in HEK293 cells before virus production. AAV1-EF1α-ChR2(H134R)-mCherry (2.0 × 10^12 gc/ml) and AAV1-Cre:GFP (5 × 10^12 gc/ml) were purchased from UNC Vector Core (Chapel Hill, NC).

### Stereotactic surgery

Mice (>4 weeks) were anesthetized with isoflurane (1-2%) and placed into a stereotaxic frame (David Kopf Instruments). For microinfusion of virus, a custom-made 30-gauge stainless injector was used to infuse 300nl of virus unilaterally at 100nl/min using a micropump (WPI UltaMicroPump). Needle was left in place for 10 min to allow for virus diffusion and prevent backflow. Following infusion, mice were allowed to recover for at least 21 days before photooxidation/EM, immunohistochemistry or electrophysiology. The following injection coordinates (in mm from bregma) were used: VTA −3.4 AP, −0.3 ML, −4.4 DV.

### Immunohistochemistry

Mice were anesthetized with ketamine (Pfizer, 10mg/kg, i.p.) and xylazine (Lloyd, 2mg/kg, i.p.). Animals were transcardially perfused with ice-cold PBS followed by 4% paraformaldehyde (PFA). Brains were incubated in 4% PFA overnight at 4°C, transferred to 30% sucrose for 48-72h until sunk and frozen in chilled isopentane. Brains were serially cut at 30 μm using a cryostat (CM3050S Leica) and collected in PBS containing 0.01% sodium azide. Immunofluorescent staining was performed as previously described (Steinkellner et al., 2019). Briefly, free-floating sections were washed three times (5 min) in PBS, blocked 1 h in PBS containing 5% normal donkey serum and 0.3% Triton X-100 (blocking buffer) followed by incubation with primary antibodies (rabbit anti-miniSOG 1:500 [Tsien Lab]; guinea pig anti-VGLUT2 1:2000 [Millipore #AB2251]; rabbit anti-TH 1:1000 [Millipore #AB152]; sheep anti-TH 1:1000 [Pel-Freeze #P60101]; rabbit Anti-Dsred 1:1000 [Clontech #632496]; mouse anti-synaptophysin 1:1000 [Synaptic Systems #101011) in blocking buffer overnight at 4°C. The following day, sections were washed three times (15 min) in PBS and incubated with secondary antibodies in blocking buffer for 2 h at room temperature. All secondary antibodies were from Jackson Immuno Research Laboratories and conjugated to either Alexa-488, Alexa-594 or Alexa-647 (5 μg/ml). Sections were rinsed three times (15 min) and mounted onto glass slides with Fluoromount-G mounting medium (Southern Biotech) ± DAPI (0.5 μg/ml). Images were captured using a Zeiss Axio Observer Epifluorescence microscope or a Leica SP5 confocal microscope.

### Electrophysiology

Adult mice (7-12 weeks) were deeply anesthetized with pentobarbital (200 mg/kg i.p.; Virbac) and perfused intracardially with 10 mL ice-cold sucrose-artificial cerebrospinal fluid (ACSF) containing (in mM): 75 sucrose, 87 NaCl, 2.5 KCl, 7 MgCl_2_, 0.5 CaCl_2_, 1.25 NaH_2_PO_4_, 25 NaHCO_3_ and continuously bubbled with carbogen (95% O_2_ – 5% CO_2_). Brains were extracted and 200 μm coronal slices were cut in sucrose-ACSF using a Leica Vibratome (vt1200). Slices were transferred to a perfusion chamber containing ACSF at 31°C (in mM): 126 NaCl, 2.5 KCl, 1.2 MgCl_2_, 2.4 CaCl_2_, 1.4 NaH_2_PO_4_, 25 NaHCO_3_, 11 glucose, continuously bubbled with carbogen. After >60 min recovery, slices were transferred to a recording chamber continuously perfused with ACSF (1-3 mL/min). Patch pipettes (3.5-6.5 MΩ) were pulled from borosilicate glass (King Precision Glass) and filled with internal recording solution containing (in mM): 120 CsCH_3_SO_3_, 20 HEPES, 0.4 EGTA, 2.8 NaCl, 5 TEA, 2.5 Mg-ATP, 0.25 Na-GTP, at pH 7.25 and 285 ± 5 mOsm.

ChR2:mCherry-labelled fibers in the NAc (ventromedial striatum) and the CPu (dorsomedial striatum) were visualized by epifluorescence and visually-guided patch recordings were made using infrared-differential interference contrast (IR-DIC) illumination (Axiocam MRm, Examiner.A1, Zeiss). ChR2 was activated by flashing blue light (473 nm; 5-ms pulse width) to trigger postsynaptic currents in whole cell through the light path of the microscope using an ultrahigh-powered light-emitting diode (LED460, Prizmatix) under computer control. Excitatory postsynaptic currents (EPSCs) were recorded in whole-cell voltage clamp with a holding potential of −65 mV (Multiclamp 700B amplifier, Axon Instruments), filtered at 2 kHz, digitized at 10 kHz (Axon Digidata 1550, Axon Instruments), and collected on-line using pClamp 10 software (Molecular Devices). Photostimuli were applied every 45 s and 10 photo-evoked currents were averaged per neuron per condition. Stock solutions were diluted 1000-fold in ACSF and bath applied at 10 μM (DNQX, Sigma) and 100 μM (APV, Sigma). Current sizes were calculated by using peak amplitude from baseline. Series resistance and capacitance were electronically compensated prior to recordings. Estimated liquid-junction potential was 12 mV and left uncorrected. Series resistance and/or leak current were monitored during recordings and cells that showed >25% change during recordings were considered unstable and discarded.

### Culture cell preparation, photooxidation and electron microscopy

Brains were extracted from postnatal day 2 Sprague Dawley rats, and cortical neurons were dissociated using papain, and plated on glass-bottom dishes (P35G-0-14-C, MatTek Corporation) coated with poly-D-lysine. Neurons were cultured in Neurobasal A medium containing 1X B27 Supplements (both from Life Technologies), 2 mM GlutaMAX (Life Technologies), 20 U/mL penicillin, and 50 mg/mL streptomycin. At day 5, neurons were co-infected with AAVDJ-Syn1-DIO-VGLUT2:mSOG and AAV-Cre-GFP, and 7 days post-infection they were fixed and processed for electron microscopy. For fixation, a pre-warmed 2% (w/v) glutaraldehyde (Electron Microscopy Sciences) in 0.1 M sodium cacodylate buffer, pH 7.4 (Ted Pella Incorporated) was added to the neurons for 5 minutes at 37°C and then plates were transferred on ice for 1 hour. Subsequently, neurons were rinsed on ice 3-5 times using chilled cacodylate buffer and treated for 30 minutes on ice in a blocking solution (50 mM glycine, 10 mM KCN, and 20 mM aminotriazole in 0.1 M sodium cacodylate buffer, pH 7.4) to reduce nonspecific background precipitation of DAB. Areas of interest were imaged and photooxidized using a Leica SPE II inverted confocal microscope outfitted with a stage chilled to 4°C. Confocal fluorescence and transmitted light images were taken with minimum exposure to identify transfected cells for correlative light microscopic imaging. For miniSOG photooxidation, DAB (3-3’-diaminobenzidine, Sigma-Aldrich catalog no. D8001-10G) was dissolved in 0.1 N HCl at a concentration of 5.4 mg ml-1 and subsequently diluted ten-fold into sodium cacodylate buffer (pH 7.4, with a final buffer concentration of 0.1 M), mixed, and passed through a 0.22 mm syringe filter before use. DAB solutions were freshly prepared on the day of photooxidation and placed on ice and protected from light before being added to cells. The samples were then illuminated through a standard FITC filter set (EX470/40, DM510, BA520) with intense light from a 150W Xenon lamp using a 63X objective lens, and oxygen was gently blown on the surface of the solution during the exposure to light. Illumination was stopped as soon as an optically-dense reaction product began to appear, as monitored by transmitted light. Neurons were then removed from the microscope, washed in ice-cold 0.1 M cacodylate buffer, and postfixed in 1% osmium tetroxide for 30 min on ice. After several washes in ice-cold double distilled water (ddH_2_O), cells were either *en bloc* stained with 2% aqueous uranyl acetate for 1 h to overnight at 4°C, or directly dehydrated in a cold graded ethanol series (20%, 50%, 70%, 90%, 100%) 3 min each on ice, then rinsed once in room temperature with 100% ethanol and embedded in Durcupan ACM resin (Electron Microscopy Sciences). In some samples the post staining step in uranyl acetate was omitted so that the specific EM signal would be generated only by the passage in osmium to add electron density to the DAB precipitates. Sections were cut with a diamond knife at a thickness of 70–90 nm for thin sections or 120 nm for EELS. Thin sections were examined using a JEOL 1200 EX operated at 80 kV.

### Mouse Brain preparation for photooxidation

Mice were anesthetized with ketamine (Pfizer, 10mg/kg, i.p.) and xylazine (Lloyd, 2mg/kg, i.p.), and then transcardially perfused with prewarmed Ringer’s solution containing xylocaine (0.2 mg/ml) and heparin (10 units/ml) for 2 minutes at 35°C, followed by 0.15 M sodium cacodylate containing 4% paraformaldehyde/0.1% glutaraldehyde/2 mM CaCl_2_ for 10 min at 35°C. The brain was then removed and placed in the same ice-cold fixative for 4 hrs. The brain was then cut into 100 μm thick slices using a vibratome and slices were incubated overnight with 0.15 M sodium cacodylate containing 4% paraformaldehyde only. The following day, the slices were washed with 0.15 M sodium cacodylate five times for 3 min on ice, and images of regions of interest were collected (both fluorescence and bright field) using a Leica SPE II inverted confocal microscope. Because VGLUT2:mSOG fluorescence is weak in thick tissue sections, we relied on co-injected ChR2:mCherry fluorescence to delineate target structures (i.e. NAc medial shell, VP and LHb) for photo-oxidation. The mouse brain slices were then post-fixed in 2.5% glutaraldehyde for 20 min on ice, then washed with 0.15 M sodium cacodylate five times for 3 min on ice. Next, the samples were incubated in 0.15 M sodium cacodylate with 10 mM KCN, 20 mM aminotriazole, 50 mM glycine, and 0.01% hydrogen peroxide for 30 min on ice. To perform photo-oxidation, freshly prepared oxygenated DAB (as indicated above) in blocking buffer was added to the dish containing the slice, and tissue was illuminated with 450–490 nm light from a 150W Xenon lamp for 30 min using a 20X objective lens. The slice was then removed from the microscope, washed in ice-cold 0.15 M cacodylate buffer, and processed for SBEM.

### Serial block-face scanning electron microscopy

Mouse brain sections were placed in 2% OsO_4_/1.5% potassium ferrocyanide in ice-cold 0.15 M cacodylate buffer for 1 h. After three 5 min washes, tissue was placed in filtered 1% thiocarbohydrazide solution for 20 min. Sections were rinsed again with ddH_2_O and then placed in a 2% OsO_4_ solution for 30 min. After this second osmium step, the sections were rinsed with ddH_2_O and left in 2% uranyl acetate aqueous solution overnight at 4°C. The next day, sections were washed five times for 2 min in ddH_2_O at room temperature and en bloc Walton’s lead aspartate staining was performed for 30 min at 60 °C. Following 5 washes for 2 min in ddH_2_O at room temperature, cells were dehydrated using a series of ice-cold graded ethanol solutions for 5 min each. Subsequently, the sections were placed into ice-cold 100% acetone for an additional 10 min, followed by a second 100% acetone step at room temperature for 10 min. The tissue was infiltrated with a solution of 50% acetone–50% Durcupan ACM epoxy resin (Electron Microscopy Sciences) for minimum 6 hrs to overnight and then placed into fresh 100% Durcupan for another day. Lastly, the sections were embedded using ACLAR and two mold-release coated glass slides and left at 60°C for 48 h.

### Image segmentation and analysis

Groundtruth segmentations of labeled and unlabeled synaptic vesicles were generated to train two deep neural network models, with CDeep3M (Haberl et al., 2018), for miniSOG labeled vesicles and unlabeled vesicles. Three subvolumes were used for training, each training set with dimensions below 1.2 × 1.2 × 2.4 μm^3^. To train the unlabeled vesicle model, one training set per region was generated from LHb and NAc and pooled for training the model. Due to a more uniform appearance of miniSOG vesicles, only one training volume from the LHb was used for training. Vesicle cross sections were extracted from the model’s predictions on the whole volume using Hough transformations and cross sections were merged across the z-plane to produce a final 3D spherical diameter. Computationally segmented vesicles were manually proof edited, facilitated by a custom written Matlab script. A separate model, trained from a variety of microscopy modes, was used to label mitochondria (Haberl et al., 2018). The pixel size for training and prediction of vesicles and mitochondria were 4.8 nm.

### Code and data availability

CDeep3M (Haberl et al., 2018) is available as open-source software on GitHub (https://github.com/CRBS/cdeep3m2) and trained deep neural network models are available at http://www.cellimagelibrary.org/cdeep3m (Haberl et al., 2020) for public use. Custom code for vesicle analysis is available at https://gitlab.nautilus.optiputer.net/madany/ves3d. SBEM data is made publicly available on the Cell Image Library (http://www.cellimagelibrary.org).

## Acknowledgements

We thank Katherine Shen, Elisabeth Soriano, and Jaskaran Shergill for help with ground truth segmentations of boutons and vesicles. This work was supported by a Schrӧdinger postdoctoral fellowship (J3656-B24) from the Austrian Science Fund (TS) and NIH grants K99AG059834 (TS), R01GM086197 (DB & SA), R01DA036612 (TSH), R01MH120685 (DB & TSH), and P41 GM103412 (MHE) for support of the National Center for Microscopy and Imaging Research.

